# Distributions of hypothalamic neuron populations co-expressing tyrosine hydroxylase and the vesicular GABA transporter in the mouse

**DOI:** 10.1101/762328

**Authors:** Kenichiro Negishi, Mikayla A. Payant, Kayla S. Schumacker, Gabor Wittmann, Rebecca M. Butler, Ronald M. Lechan, Harry W. M. Steinbusch, Arshad M. Khan, Melissa J. Chee

**Affiliations:** UTEP Systems Neuroscience Laboratory, Department of Biological Sciences, and Border Biomedical Research Center, University of Texas at El Paso, El Paso, TX 79968, USA; Department of Neuroscience, Carleton University, Ottawa, ON, K1S 5B6, Canada; Department of Medicine, Division of Endocrinology, Diabetes and Metabolism, Tufts Medical Center, Boston, MA 02111, USA; Department of Psychiatry and Neuropsychology, School for Mental Health and Neuroscience, Maastricht University, Maastricht, Netherlands

**Keywords:** zona incerta, dopamine, hypothalamus, GABA, tyrosine hydroxylase, vGAT, catecholamine

## Abstract

The hypothalamus contains catecholaminergic neurons marked by the expression of tyrosine hydroxylase (TH). As multiple chemical messengers coexist in each neuron, we determined if hypothalamic TH-immunoreactive (ir) neurons express glutamate or GABA. We used Cre/loxP recombination to express enhanced GFP fluorescence (EGFP) in neurons that express the vesicular glutamate (vGLUT2) or GABA transporter (vGAT), then determined TH immunoreactivity in glutamatergic or GABAergic neurons, respectively. EGFP-positive vGLUT2 neurons were not TH-ir. However, discrete TH-ir signals colocalized with EGFP-positive vGAT neurons, which we validated by *in situ* hybridization for *Vgat* mRNA. In order to contextualize the observed pattern of TH+EGFP colocalization in vGAT neurons, we first performed Nissl-based parcellation and plane-of-section analysis, and then mapped the distribution of TH-ir vGAT neurons onto atlas templates from the *Allen Reference Atlas (ARA)* of the mouse brain. TH-ir vGAT neurons were distributed throughout the rostrocaudal extent of the hypothalamus. Within the *ARA* ontology of gray matter regions, TH-ir neurons localized primarily to the periventricular hypothalamic zone, periventricular hypothalamic region, and lateral hypothalamic zone. There was a very strong presence of EGFP fluorescence in TH-ir neurons across all brain regions, but the most striking colocalization was found in the zona incerta (ZI) – a region assigned to the hypothalamus in the *ARA* – where every TH-ir neuron was EGFP-positive. Neurochemical characterization of these ZI neurons revealed that they display immunoreactivity for dopamine but not dopamine β-hydroxylase. In aggregate, these findings indicate the existence of a novel hypothalamic population in the mouse that may signal through the release of GABA and/or dopamine.

## 1. Introduction

The prevailing view of neurotransmission is that multiple chemical messengers can coexist, in various combinations, within single neurons and in some cases can even be co-stored in synaptic vesicles (Hökfelt et al., 1986). These chemical messengers may be neurotransmitters like glutamate and GABA that rapidly initiate and terminate discrete synaptic events, or neuromodulators like neuropeptides and catecholamines that can have varied release probabilities, time courses of release, or long-range target sites. In fact, virtually all neuropeptidergic neurons also harbor a neurotransmitter, as exemplified by brain structures in the hypothalamus. For example, neurons in the arcuate hypothalamic nucleus coexpress the neuropeptides Agouti-related protein and neuropeptide Y (Broberger et al., 1998a; Broberger et al., 1998b; Hahn et al., 1998) as well as the neurotransmitter GABA (Tong et al., 2008).

Colocalized neurotransmitters and neuromodulators may work in tandem or opposition to prolong or suppress neuronal functions (van den Pol, 2012). For instance, neurons in the lateral hypothalamus that express either melanin-concentrating hormone (MCH) or hypocretin/orexin (H/O) have transcripts encoding the machinery for vesicular storage of glutamate (Mickelsen et al., 2017). Both of these neuronal populations can release glutamate and produce transient excitatory glutamatergic events (Schöne et al., 2014; Chee et al., 2015), but MCH inhibits (Wu et al., 2009; Sears et al., 2010) whereas H/O stimulates neuronal activity (Schöne et al., 2014). Additionally, these coexpressed messengers may also serve distinct functions. For example, MCH, but not glutamate from MCH neurons, promotes rapid eye movement sleep (Naganuma et al., 2019), while glutamate has been shown to encode the nutritive value of sugars (Schneeberger et al., 2018).

The colocalization between neuropeptides and catecholamines is also well-studied. The catecholamines are a major class of monoamine messengers that include dopamine, norepinephrine, and epinephrine. Their presence in neurons is indirectly indicated by the detection of immunoreactivity for tyrosine hydroxylase (TH; EC 1.14.16.2), which mediates the rate-limiting step in catecholamine biosynthesis (Udenfriend & Wyngaarden, 1956; Nagatsu et al., 1964). Discrete populations of catecholamine-containing neurons, and specifically TH-immunoreactive (ir) neurons, have been reported in the hypothalamus (Björklund & Nobin, 1973; Hökfelt et al., 1976; Björklund & Lindvall, 1984), and they may also colocalize with additional neurotransmitters and/or neuropeptides. Hypothalamic TH-ir neurons are primarily found within the zona incerta (ZI) and periventricular parts of the hypothalamus (Hökfelt et al., 1976; Ruggiero et al., 1984), which is composed of neurochemically and functionally diverse gray matter regions. For example, hypothalamic dopamine neurons may colocalize with galanin and neurotensin as well as with markers for GABA synthesis (Everitt et al., 1986). Furthermore, recent work has demonstrated that dopaminergic neurons in the arcuate nucleus corelease GABA (Zhang & van den Pol, 2015), and that both neurotransmitters regulate feeding in response to circulating metabolic signals (Zhang & van den Pol, 2016).

Single-cell RNA sequencing has revealed that neurochemically defined populations, such as hypothalamic TH-expressing neurons, can be further sorted into transcriptionally distinct subgroups (Romanov et al., 2017). Gene expression-based cluster analysis is a powerful tool for identifying subpopulations within seemingly homogeneous cell groups (Romanov et al., 2017). However, it is necessary for these transcriptomic studies to be supported and framed by accurate structural information (Crosetto et al., 2015; Khan et al., 2018). Consideration of basic properties, such as morphology, connectivity, and spatial distributions can reveal groupings (Bota & Swanson, 2007), which would likely remain undetected by high-throughput transcriptomic analyses alone. Indeed, for the hypothalamus, most published transcriptomic and proteomic analyses have ignored cytoarchitectonic boundary conditions of the diverse gray matter regions in this structure, opting instead to report gene expression patterns from the whole hypothalamus (Khan et al., 2018).

We therefore examined the spatial distributions of hypothalamic TH neurons and quantified the extent of their colocalization with other neurotransmitters or neuropeptides. We first determined whether hypothalamic TH-ir neurons may be either GABAergic or glutamatergic. By combining multi-label immunohistochemistry with cytoarchitectural analysis to obtain high spatial resolution maps of TH-ir neuronal subpopulations, we found that hypothalamic TH-ir neurons were predominantly GABAergic, and that this colocalization was especially prevalent within a specific region of the rostral ZI. We further characterized the neurochemical identity of ZI GABAergic TH-ir neurons and showed that these are dopaminergic neurons, but they do not colocalize with known neuropeptides in the ZI region.

## 2. Materials & Methods

All animal care and experimental procedures were completed in accordance with the guidelines and approval of the Animal Care Committee at Carleton University. Mice were housed in ambient temperature (22–24°C) with a 12:12 light dark cycle with ad libitum access to water and standard mouse chow (Teklad Global Diets 2014, Envigo, Mississauga, Canada).

### 2.1: Generation of *Vgat-cre;L10-Egfp* and *Vglut2-cre;L10-Egfp* mice

To visualize GABAergic and glutamatergic neurons, respectively, we labeled neurons expressing the vesicular GABA transporter (vGAT; *Slc32a1*) or vesicular glutamate transporter 2 (vGlut2; *Scl17a6*) with enhanced GFP fluorescence by crossing a *Vgat-ires-cre* (Stock 028862, Jackson Laboratory, Bar Harbor, ME) or *Vglut2-ires-cre* mouse (Stock No. 16963, Jackson Laboratory) with a Cre-dependent *lox-STOP-lox-L10-Egfp* reporter mouse (Krashes et al., 2014), kindly provided by Dr. B. B. Lowell (Beth Israel Deaconess Medical Center, Boston, MA) to produce *Vgat-cre;L10-Egfp* and *Vglut2-cre;L10-Egfp* mice.

### 2.2: Antibody characterization

Table 1 lists the following primary antibodies we used for immunohistochemistry (IHC).

**Table 1.** Details of the primary antibodies used in this study

| Antibody | Immunogen | Clonality | Isotype | Source | Cat. # | Lot # | RRID | Titer |
| --- | --- | --- | --- | --- | --- | --- | --- | --- |
| sheep anti-DIG | isolated from sheep immunized with DIG steroid molecule | polyclonal | IgG | Roche | 11207733910 | 28557000 | AB_514500 | 1:100 |
| goat anti-dopamine | dopamine-glutaraldehyde-thyroglobulin conjugate | polyclonal | IgG | Dr. H. Steinbusch, Maastricht University | — | — | na | 1:2,000 |
| rabbit anti-DBH | purified enzyme from bovine adrenal medulla | polyclonal | IgG | Immunostar | 22806 | 1233001 | AB_572229 | 1:5,000 |
| chicken anti-GFP | His-tagged GFP isolated from *Aequorea victoria* | polyclonal | IgY | Millipore | 06-896 | 3022861 | AB_11214044 | 1:2,000 |
| rabbit anti-GFP | isolated from *Aequorea victoria* | polyclonal | IgG | Invitrogen | A6455 | 52630A | AB_221570 | 1:1,000 |
| rabbit anti-MCH | full-length MCH peptide | polyclonal | IgG | Dr. E. Maratos-Flier, Beth Israel Deaconess Medical Center | — | — | AB_2314774 | 1:5,000 |
| goat anti-orexin-A | C-terminus of human orexin peptide | polyclonal | IgG | Santa Cruz | sc-8070 | J2414 | AB_653610 | 1:4,000 |
| mouse anti-TH | purified from N-terminus in PC12 cells | monoclonal | IgG | Millipore | MAB318 | 2677893 | AB_2201528 | 1:2,000 |

DIG, digoxigenin; DBH, dopamine beta-hydroxylase; GFP, green fluorescent protein; MCH, melanin-concentrating hormone; na, not available; TH, tyrosine hydroxylase; —, not applicable

#### Sheep anti-digoxigenin (DIG) antibody

When compared to tissue incubated with a DIG-labeled riboprobe followed with the anti-DIG antibody, no hybridization signals were detected when the tissue was incubated with the anti-DIG antibody only (Vazdarjanova et al., 2006).

#### Goat anti-dopamine antibody

Specificity was determined through a gelatin model system and nitrocellulose sheets. The anti-dopamine antibody showed immunoreactivity to even low concentrations of dopamine, with cross-reactivity of less than 10% to noradrenaline, less than 1% to other monoamines, and low levels of cross-reactivity to L-DOPA at higher concentrations (Steinbusch et al., 1991).

#### Rabbit anti-dopamine β-hydroxylase

Specificity was determined by the absence of staining if the antibody was first preadsorbed by the DBH peptide (Sockman & Salvante, 2008). The distribution of DBH immunoreactivity was confirmed in the locus coeruleus (data not shown), a region known to express DBH, as previously shown (Yamaguchi et al., 2018). This antibody has also been reported to reveal a distribution of hindbrain DBH-immunoreactive neurons that is similar to that revealed by a DBH antibody of different origin (Halliday & McLachlan, 1991).

#### Chicken anti-green fluorescent protein (GFP) antibody

Wild type brain tissue does not endogenously express the GFP transgene and incubating wild type brain tissue with this antibody did not produce any GFP-ir signals (data not shown).

#### Rabbit anti-GFP antibody

Specificity was determined by the absence of GFP immunoreactivity in the brain of wild type *Drosophila melanogaster*, which does not endogenously produce endogenous GFP molecules (Busch et al., 2009).

#### Rabbit anti-melanin-concentrating hormone (MCH) antibody

The polyclonal anti-MCH antibody was made and generously provided by Dr. E. Maratos-Flier (Beth Israel Deaconess Medical Center, Boston, MA). Specificity was determined by the lack of MCH immunoreactivity after preadsorption with MCH peptide (Elias et al., 1998) or after application to brain tissue from MCH knockout mice (Chee et al., 2013).

#### Goat anti-orexin-A antibody

Specificity was demonstrated by preadsorption with orexin peptide, which abolished all specific staining shown with this antibody (Florenzano et al., 2006).

#### Mouse anti-tyrosine hydroxylase (TH) antibody

Knockdown of *th1* gene that encodes Th in the central nervous system of the zebrafish abolished detectable immunoreactivity from this antibody (Kuscha et al., 2012). We did not observe any TH-ir signals when this antibody was added to either wild type or *Vgat-cre;L10-Egfp* brain tissue (data not shown).

All secondary antibodies (Table 2) were raised in donkey against the species of the conjugated primary antisera (mouse, goat, rabbit, or sheep).

**Table 2.** Details of the secondary antibodies used in this study

| Antibody | Isotype | Source | Cat. # | Lot # | RRID | Titer |
| --- | --- | --- | --- | --- | --- | --- |
| AffiniPure donkey anti-chicken Alexa Fluor 488 | IgY (H+L) | Jackson ImmunoResearch | 703-545-155 | 139424 | AB_2340375 | 1:500 |
| AffiniPure donkey anti-goat Alexa Fluor 488 | IgG (H+L) | Jackson ImmunoResearch | 705-545-147 | 103943 | AB_2336933 | 1:500 |
| AffiniPure donkey anti-rabbit Alexa Fluor 488 | IgG (H+L) | Jackson ImmunoResearch | 711-545-152 | 125719 | AB_2313584 | 1:200 |
| Donkey anti-rabbit Alexa Fluor 488 | IgG (H+L) | Thermo Fisher Scientific | A-21206 | 19107512 | AB_141708 | 1:500 |
| Donkey anti-mouse Alexa Fluor 568 | IgG (H+L) | Thermo Fisher Scientific | A-10037 | 1827879 | AB_2534013 | 1:500 |
| Donkey anti-goat Alexa Fluor 647 | IgG (H+L) | Thermo Fisher Scientific | A-21447 | 1977345 | AB_2535864 | 1:500 |
| Donkey anti-rabbit Alexa Fluor 647 | IgG (H+L) | Thermo Fisher Scientific | A-31573 | 1903516 | AB_2536183 | 1:300 |

### 2.3: Immunohistochemistry (IHC)

#### 2.3.1: Tissue processing

Male and female *Vgat-cre;L10-Egfp*, *Vglut2-cre;L10-Egfp*, and wild type mice on a mixed C57BL/6, FVB, and 129S6 background (8–15 weeks old) were anesthetized with an intraperitoneal (ip) injection of urethane (1.6 g/kg) and transcardially perfused with cold (4°C) 0.9% saline followed by 10% neutral buffered formalin (4°C), unless indicated otherwise. The brain was removed from the skull, post-fixed with 10% formalin for 24 hours at 4°C and then cryoprotected in phosphate-buffered saline (pH 7.4, 0.01 M) containing 20% sucrose and 0.05% sodium azide (4°C). Brains were cut into four or five series of 30 μm-thick coronal-plane sections using a freezing microtome (Leica SM2000R, Nussloch, Germany) and stored in an antifreeze solution containing 50% formalin, 20% glycerol, and 30% ethylene glycol.

To examine dopamine immunoreactivity, *Vgat-cre;L10-Egfp* brains were perfused with cold (4°C) saline and then 1% glutaraldehyde:9% formalin mixture (4°C). The brains were post-fixed in the perfusion solution for 30 minutes and cryoprotected with 30% sucrose solution for 24 hours (4°C). Brains were cut into five series of 30 μm-thick sections and collected into a PBS-azide solution containing 1% formalin. All aforementioned perfusion and incubation solutions contained 1% sodium metabisulfite to prevent the oxidation of dopamine.

#### 2.3.2: Procedure

Single- and dual-label IHC were completed as previously described (Chee et al., 2013), unless indicated otherwise, using the antibodies and dilution combinations listed in Table 1 and Table 2. In brief, brain tissue sections were washed with six 5-minute PBS exchanges and then blocked for 2 hours at room temperature (RT, 20–21°C) with 3% normal donkey serum (Jackson ImmunoResearch Laboratories, Inc., West Grove, PA) dissolved in PBS-azide containing 0.25% Triton X-100 (PBT-azide). Unless indicated otherwise, primary antibodies were concurrently added to the blocking serum and incubated with the brain sections overnight (16–18 hours, RT). After washing with six 5-minute PBS rinses, the appropriate secondary antibodies were diluted in PBT containing 3% NDS and applied to the brain tissue sections for 2 hours (RT). The sections were rinsed with three 10-minute PBS washes before being mounted on SuperFrost Plus glass microscope slides (Fisher Scientific, Pittsburgh, PA) and coverslipped (#1.5 thickness) using ProLong Gold antifade reagent containing DAPI (Fisher Scientific, Pittsburgh, PA).

To examine dopamine immunoreactivity, *Vgat-cre;L10-Egfp* brain tissues were incubated in PBS containing 0.5% sodium borohydride for 30 minutes at RT, followed by six 5-minute washes in PBS. The brain sections were then blocked with 3% NDS in PBT-azide for 2 hours prior to the simultaneous overnight-incubation (RT) of primary antibodies against GFP (anti-chicken) and TH, then washed with six 5-minute exchanges in PBS, and incubated with the corresponding Alexa Fluor 488- and Alexa 568-conjugated secondary antibodies in 3% NDS for 2 hours. After three 10-minute PBS rinses of the tissue sections, we diluted the dopamine primary antibody into a 3% NDS, PBT-azide solution and incubated the sections in this mixture overnight at RT. The sections were rinsed with PBS six times for 5 minutes each before incubating them with the corresponding Alexa Fluor 647-conjugated secondary antibody in 3% NDS-PBT. Following three 10-minute washes in PBS, the brain sections were mounted and coverslipped. All solutions used to process the visualization of dopamine immunoreactivity contained 1% sodium metabisulfite to prevent the oxidation of dopamine.

### 2.4: Dual fluorescence *in situ* hybridization (fISH) and IHC

#### 2.4.1: Tissue preparation for fISH

*Vgat-cre;L10-Gfp* mice were anesthetized with urethane (1.6 g/kg, ip), and their brains were rapidly removed and snap-frozen on powdered dry ice. The brains were thawed to –16°C and sliced into ten series of 16 μm-thick coronal sections through the entire hypothalamus using a cryostat (CM3050 S, Leica Microsystems, Buffalo Grove, IL). Each section was thaw-mounted on Superfrost Plus slides (Fisher Scientific), air-dried at RT, and then stored at –80°C until processed for fISH.

#### 2.4.2: fISH and IHC procedure

The antisense vGAT riboprobe corresponds to nucleotide 875–1,416 of mouse *Slc32a1* mRNA (*Vgat*; NM_009508.2) (Agostinelli et al., 2017). Using a plasmid linearized with SacI (New England Biolabs, Beverly, MA), it was transcribed with T7 polymerase (Promega, Madison, WI) in the presence of digoxigenin (DIG)-conjugated UTP (Roche Diagnostics, Mannheim, Germany).

fISH for *Vgat* mRNA was performed according to the protocol previously described for fresh frozen sections (Wittmann et al., 2013). In brief, prior to hybridization, the sections were fixed with 4% paraformaldehyde for 20 minutes and rinsed with two 3-minute PBS washes; acetylated with 0.25% acetic anhydride in 0.1 M triethanolamine and rinsed with two PBS washes; then dehydrated in an ethanol gradient starting from 80%, 95%, to 100% for 1 minute each, chloroform for 10 minutes, and again in 100% and 95% ethanol for 1 minute each. The riboprobe (650 pg/μl) was mixed with hybridization buffer containing 50% formamide, 2× sodium citrate buffer (SSC), 1× Denhardt’s solution (MilliporeSigma, Burlington, MA), 0.25 M Tris buffer (pH 8.0), 10% dextran sulfate, 3.5% dithiothreitol, 265 μg/ml denatured salmon sperm DNA (MilliporeSigma), and applied directly to each slide under a plastic Fisherbrand coverslip (Fisher Scientific). Hybridization occurred overnight at 56°C in humidity chambers. Following hybridization, we removed the coverslip and the sections were washed in 1× SSC for 15 minutes; treated with 25 μg/ml RNase A (MilliporeSigma) dissolved in 0.5 M NaCl, 10 mM Tris, 1 mM EDTA (pH 8.0) for 1 hour (37°C); sequentially washed at 65°C with 1× SSC for 15 minutes, 0.5× SSC for 15 minutes, and 0.1× SSC for 1 hour. Subsequently, the sections were treated with PBS containing 0.5% Triton X-100 and 0.5% H_2_O_2_ (pH 7.4) for 15 minutes, rinsed in three 10-minute PBS washes, then immersed in 0.1 M maleate buffer (pH 7.5) for 10 minutes before blocking in 1% Blocking Reagent (Roche) for 10 minutes. The sections were incubated in horseradish peroxidase-conjugated, sheep anti-DIG antibody in blocking reagent and enclosed over sections with a CoverWell incubation chamber (Grace Bio-Labs Inc., Bend, OR) overnight at 4°C. After rinsing with three 10-minute PBS washes, the hybridization signal was amplified with a TSA Plus Biotin Kit (Perkin Elmer, Waltham, MA), by diluting the TSA Plus biotin reagent (1:400) in 0.05 M Tris (pH 7.6) containing 0.01% H_2_O_2_, for 30 minutes. The sections were incubated with streptavidin-conjugated Alexa Fluor 555 (1:500; S32355, Invitrogen; RRID: AB_2571525) in 1% Blocking Reagent for 2 hours (RT) to label the *Vgat* mRNA hybridization signal.

Following fISH, we processed the sections for immunofluorescence by incubating them with a GFP (rabbit) antibody, rinsing with three 10-minute PBS washes, and labeling with an Alexa Fluor 488-conjugated anti-rabbit secondary (Table 2). Finally, the sections were then washed in PBS and coverslipped with Vectashield mounting medium containing DAPI (H-1200, Vector Laboratories, Burlingame, CA).

### 2.5: Microscopy

All images were exported as TIFF files into Adobe Illustrator CS4 (Adobe Systems Inc., San Jose, CA) for assembly into multi-panel figures and to add text labels and arrowheads.

#### 2.5.1: Epifluorescence imaging

Images showing the colocalization between *Vgat* mRNA and GFP-ir in *Vgat-cre;L10-Egfp* tissue were captured using a Zeiss Axioplan 2 microscope (Carl Zeiss Inc., Göttingen, Germany) equipped with a RT SPOT digital camera (Diagnostic Instruments, Sterling Heights, MI). Adobe Photoshop CS4 (Adobe) was used to create composite images and to modify brightness or contrast to increase the visibility of lower level signals.

Large field-of-view images of whole brain sections were acquired with a fully motorized Nikon Ti2 inverted microscope (Nikon Instruments Inc., Missisauga, Canada) mounted with a Prime 95B CMOS camera (Photometrics, Tucson, AZ) using a CF160 Plan Apochromat ×20 objective lens (0.75 numerical aperture). Images were stitched using NIS Elements software (Nikon), which was also used to adjust image brightness and contrast.

#### 2.5.2: Confocal imaging

High-magnification confocal confocal stacks (2048 × 2048 pixels) involving two- or three-color fluorescence channels were visualized using a Nikon C2 confocal microscope fitted with Plan Apochromat ×20 (0.75 numerical aperture) or ×40 objective lenses (0.95 numerical aperture) and acquired using NIS Elements software (Nikon). The excitation light was provided by 488-nm, 561-nm, and 640-nm wavelength lasers for the visualization of Alexa Fluor 488, Alexa Fluor 568, and Alexa Fluor 647, respectively. We used NIS Elements to stitch overlapping frames, flatten confocal stacks by maximum intensity projection, and adjust the brightness or contrast of flattened images. Two-color images were coded in green (Alexa Fluor 488) and pseudo-colored magenta (Alexa Fluor 568). Three-color images were coded in green (Alexa Fluor 488), red (Alexa Fluor 568), and pseudo-colored light blue (Alexa Fluor 647).

### 2.6: Nissl-based parcellations, plane-of-section analysis, and atlas-based mapping

An Olympus BX53 microscope (Olympus Corporation, Tokyo, Japan) was used to examine Nissl-stained tissue sections. Photomicrographs were produced with an Olympus DP74 color camera powered by CellSens Dimension software (Version 2.3). Contrast enhancements and brightness adjustments were made with Photoshop (Version CS6; Adobe) before analyses. Photomicrographs of Nissl-stained tissue were imported to Adobe Illustrator (Version CC 2014, Adobe) and regional boundaries were drawn on a separate data layer. Line parcellations were made using the nomenclature and boundary definitions of the *Allen Reference Atlas* (*ARA*; Dong, 2008). To the extent that was applicable, cytoarchitectural criteria outlined in the rat brain atlas of Swanson (2018) were also used to clarify delineations. Parcellations were then carefully superimposed on corresponding fluorescence images to precisely reveal the spatial distributions of TH- and vGAT-ir neurons. We photographed fluorescently-immunostained tissue sections using a Zeiss Axio Imager M.2 microscope (Carl Zeiss Corporation, Thornwood, NY) using ×10 (0.3 numerical aperture) and ×20 (0.8 numerical aperture) Plan Apochromat objective lenses. An EXi Blue monochrome camera (Teledyne Qimaging, Inc., Surrey, British Columbia, Canada) was used to capture multi-channel fluorescence images and a motorized stage controlled by Volocity Software (Version 6.1.1; Quorum Technologies, Inc., Puslinch, Ontario, Canada) aided in generating stitched mosaic images. Image files were then exported in TIFF format to allow further processing using Adobe Photoshop.

Using the parcellations that were superimposed onto the corresponding epifluorescence image, cells that existed within the parcellated boundaries were marked using the *Blob Brush Tool* in Adobe Illustrator. A red circle represented a TH-positive cell, and a blue circle represented a TH- and vGAT-positive cell. Plane-of-section analysis (Zséli et al., 2016) was performed to assign each photographed tissue section or portion thereof to the appropriate *ARA* reference atlas level(s). The cell representations were then superimposed onto the corresponding *ARA* reference atlas template (Dong, 2008) for each atlas level, thus creating atlas-based maps of the tissue of interest.

### 2.7: Cell counting

The numbers of cells counted from the procedures below are semi-quantitative measures meant to provide data for relative comparisons rather than absolute cell numbers within the hypothalamus.

#### 2.7.1: Quantification of Vgat hybridization signals

We acquired stitched epifluorescence images for whole hypothalamic sections using a ×10 objective on a fully motorized Olympus BX61VS microscope running VS-ASW-FL software (Olympus). We viewed the images offline using OlyVIA software (Olympus) in order to quantify the colocalization of *Vgat* hybridization signals, which expressed red fluorescence, in GFP-ir neurons from *Vgat-cre;L10-Egfp* brain tissue. Cells were counted using a grid comprising 1-inch × 1-inch squares that was printed on a transparency. This grid was overlaid onto the magnified image on the computer screen so that each grid encompassed a 100-μm^2^ area of the brain. We counted from one hemisphere and placed the first counting square starting at the ventral edge of the base of the brain, *i.e.*, at the median eminence, then working dorsally and away from the third ventricle until the cerebral peduncle. In order to avoid double-counting, cells that landed on the gridlines were not included in the count. The percentage of colocalization was quantified by determining the number of *Vgat* mRNA neurons relative to the number of GFP-ir neurons.

#### 2.7.2: Quantification of EGFP fluorescence and TH immunoreactivity

We acquired stitched epifluorescence images using a ×10 objective (0.3 numerical aperture) on an Olympus BX53 microscope mounted with the Olympus DP74 camera, and cell counts were determined using Adobe Illustrator CS5 (Adobe). We counted all neurons displaying TH immunoreactivity, which exhibited red fluorescence, and determined the proportion of these neurons that also displayed EGFP fluorescence. The percentage of colocalization was calculated with respect to the parcellation-based cytoarchitectural boundaries and only the neurons that fell within the defined boundaries were counted, unless indicated otherwise in the Results.

### 2.8: Statistics

Line of best-fit for linear regressions was determined using Prism 6.07 (GraphPad Software Inc., San Diego, CA). All frequency distribution histograms were generated with Prism. All data are expressed as the mean ± standard error of the mean (SEM).

## 3. Results

### 3.1: Catecholaminergic neurons in the hypothalamus may be GABAergic

Catecholaminergic neurons are distributed throughout the hypothalamus (Dahlström & Fuxe, 1964; Hökfelt et al., 1976; Björklund & Lindvall, 1984; Ruggiero et al., 1984), and we determined if they may coexpress the neurotransmitter glutamate or GABA, which can be marked by their expression of vGLUT2 or vGAT, respectively. We therefore examined if TH-ir neurons in *Vglut2-cre;L10-Egfp* and *Vgat-cre;L10-Egfp* mouse brain tissue, respectively, also express native EGFP fluorescence (EGFP-f). Interestingly, we did not observe any TH-ir EGFP-f neurons from the hypothalamus of *Vglut2-cre;L10-Egfp* mice (Fig. 1); in contrast, colocalization between TH-ir and EGFP-f occurred in *Vgat-cre;L10-Egfp* brain tissue (Fig. 2). These observations suggested that hypothalamic TH-ir neurons may be GABAergic but not glutamatergic.

**Figure 1.**
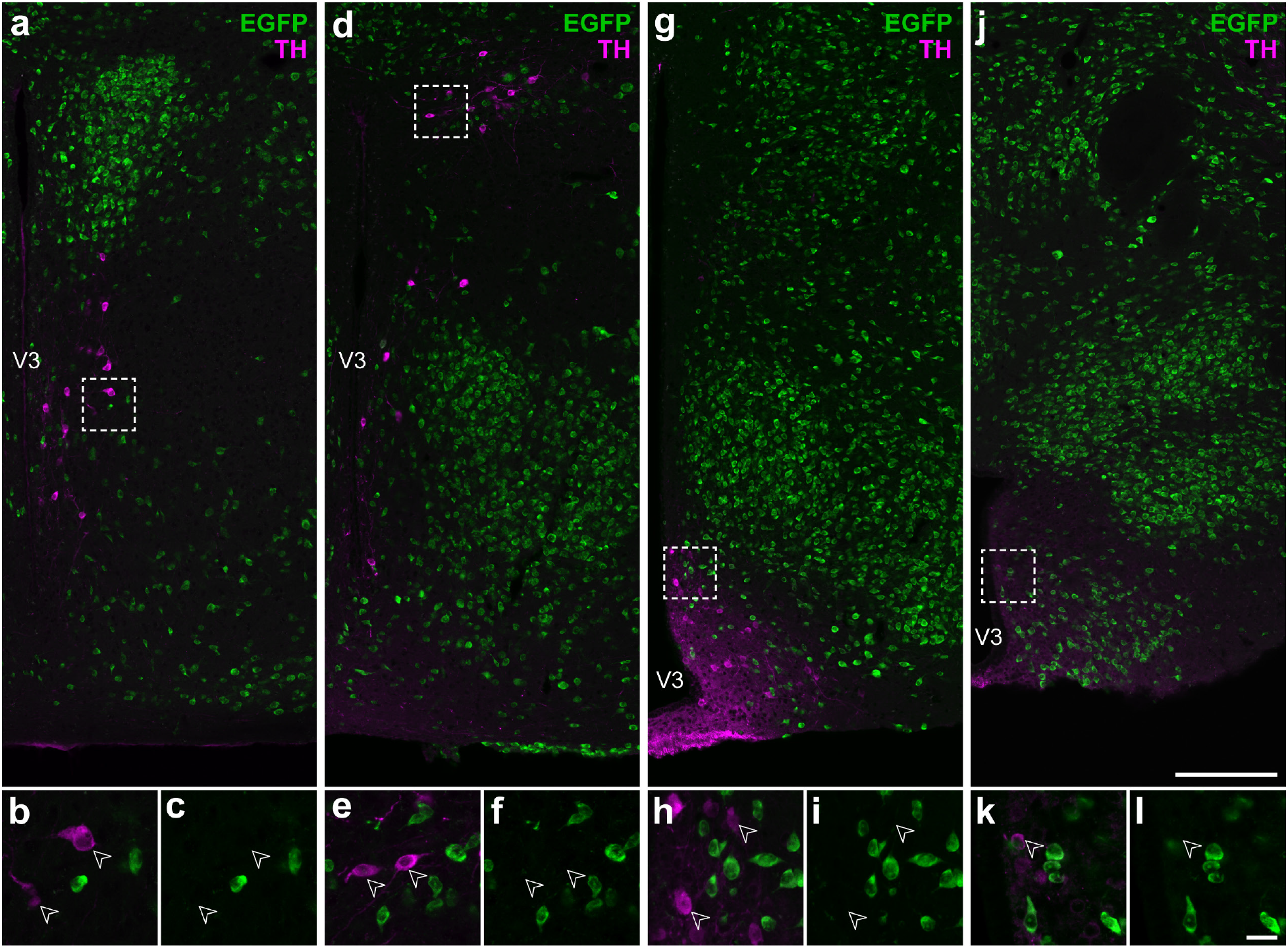
Hypothalamic catecholamine neurons are not glutamatergic. Representative stitched confocal photomicrographs showing the medial hypothalamic zone within 600 μm of the third ventricle (V3) in *Vglut2-cre;L10-Egfp* mice at the inferred anteroposterior positions (in mm) from Bregma: –0.5 **(a)**, –1.3 **(d)**, –1.6 **(g)**, and –2.2 **(j)**. Merged-channel confocal photomicrographs **(b, e, h, k)** from the respective outlined areas **(a, d, g, j)** indicate that TH-ir neurons (*open white arrowheads*) do not express native EGFP fluorescence (*green*; **c, f, i, l**). Scale bars: 200 μm in **j** also applies to **a**, **d**, **g**; 20 μm in **l** also applies to **b, c, e, f, h, i, k**.

**Figure 2.**
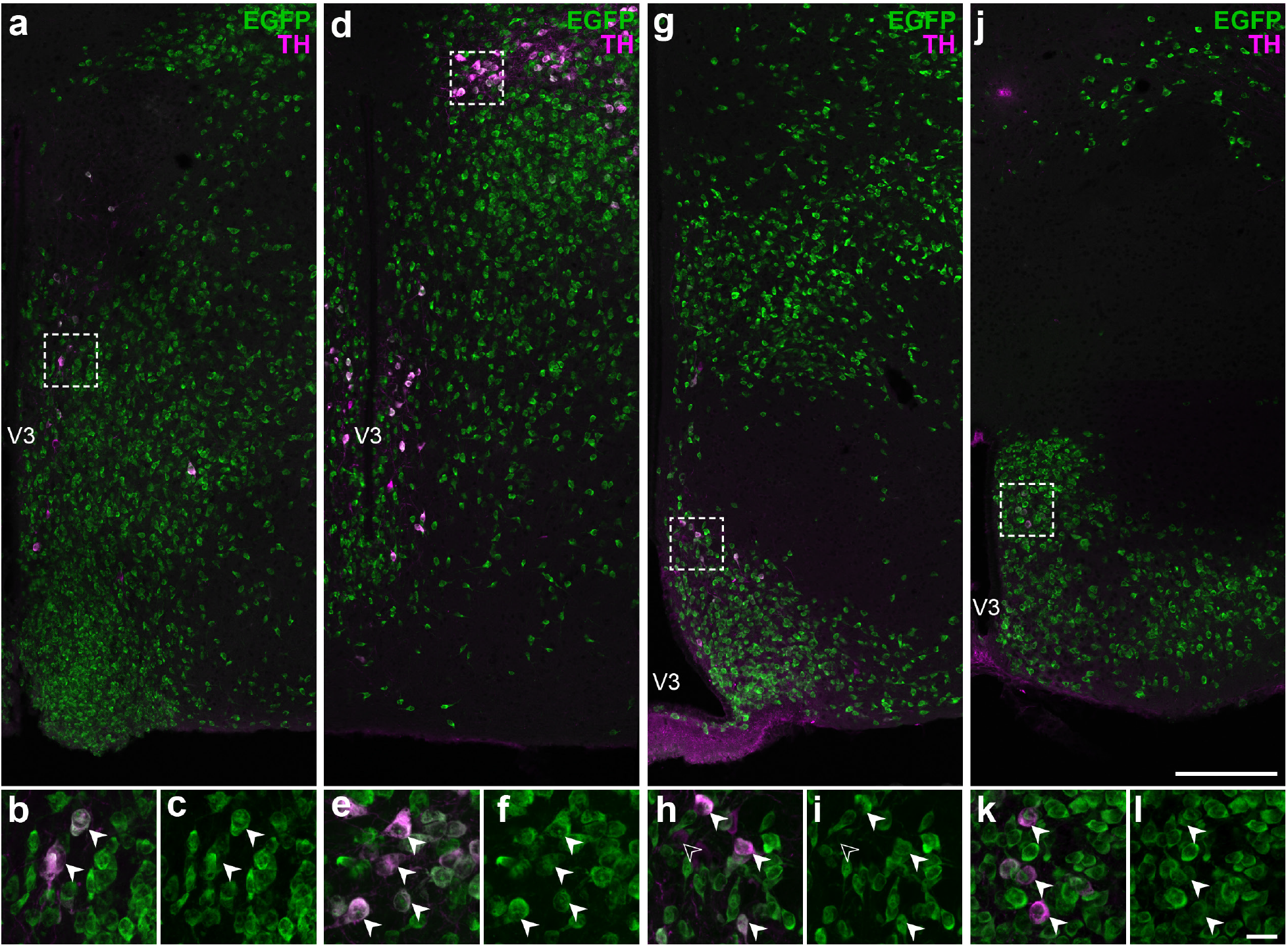
Some hypothalamic catecholamine neurons are GABAergic. Representative stitched confocal photomicrographs showing the medial hypothalamic zone within 600 μm of the third ventricle (V3) in *Vgat-cre;L10-Egfp* mice at the inferred anteroposterior positions (in mm) from Bregma: –0.5 **(a)**, –1.3 **(d)**, –1.6 **(g)**, and –2.2 **(j)**. Merged-channel confocal photomicrographs **(b, e, h, k)** from the respective outlined areas in **(a, d, g, j)** indicate that the vast majority of TH-ir neurons (*filled white arrowheads*) coexpress native EGFP fluorescence (*green*; **c, f, i, l**). *Open white arrowheads* **(h, i)** mark TH-ir neurons that do not coexpress EGFP fluorescence. Scale bars: 200 μm in **j** also applies to **a, d, g**; 20 μm in **l** also applies to **b, c, e, f, h, i, k**.

We evaluated the efficacy and specificity of EGFP-f to indicate vGAT-expressing neurons in *Vgat-cre;L10-Egfp* mice by assessing the colocalization of *Vgat* mRNA hybridization (*Vgat*-ISH) in GFP-ir neurons (Fig. 3). We counted GFP-ir from one hemisphere of the hypothalamus and found that more than 99% of GFP-ir neurons (18,697 out of 18,805) expressed *Vgat*-ISH signals. Thus EGFP-positive neurons have nearly complete one-to-one correspondence with *Vgat*-ISH signal in the hypothalamus. Furthermore, less than 1% of the *Vgat*-ISH neurons counted (138 out of 18,835) were not GFP-ir in the hypothalamus, so the likelihood of under-reporting a *Vgat*-positive neuron was minimal. These findings validated the use of the *Vgat-cre;L10-Egfp* mouse model for unambiguous detection of GABAergic vGAT-positive hypothalamic neurons by EGFP-f.

**Figure 3.**
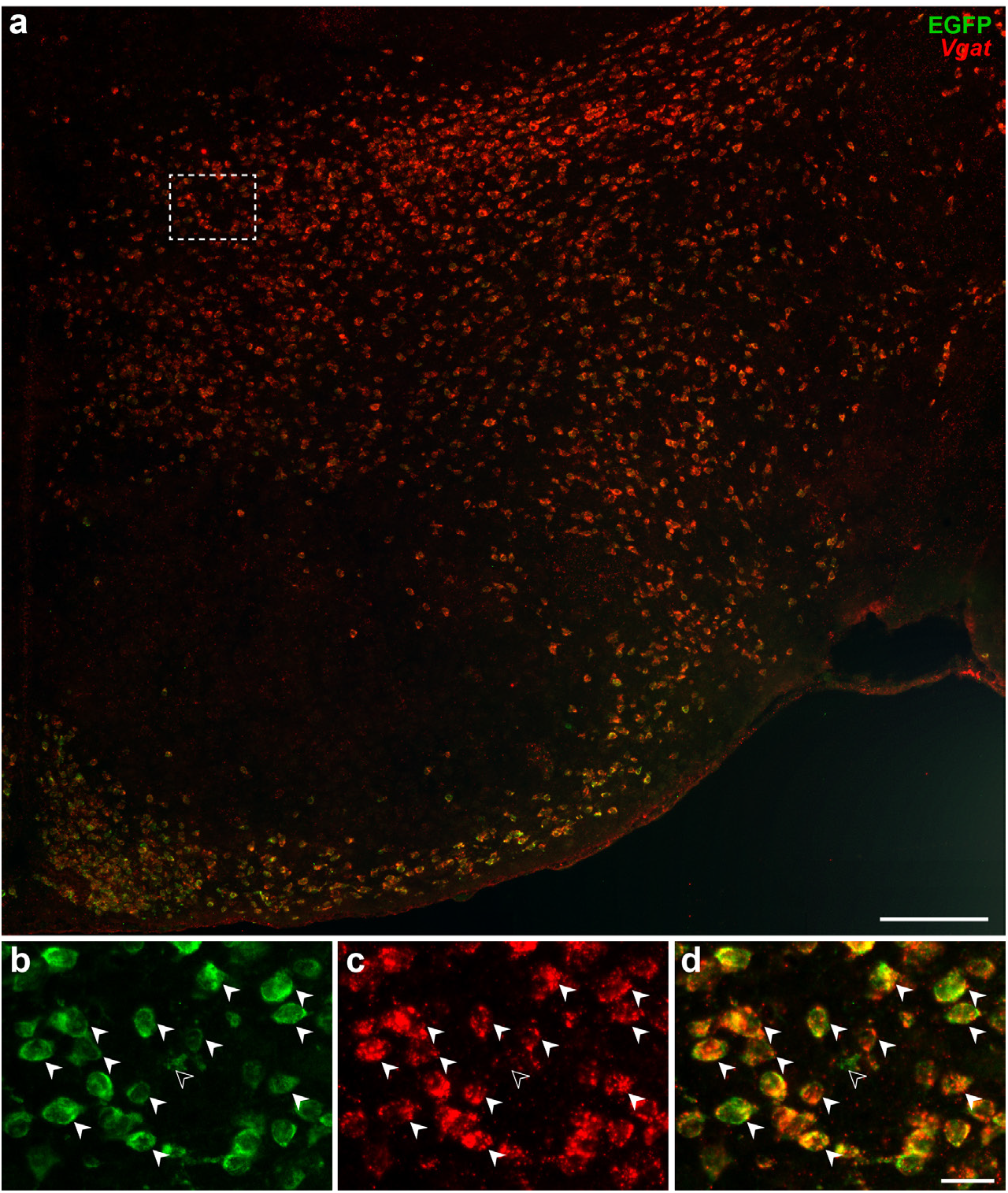
Visualization of GABAergic vGAT neurons by GFP fluorescence. Epifluorescence photomicrograph showing *vGAT* mRNA hybridization signals in GFP-ir neurons from *Vgat-cre;L10-Egfp* mice. High magnification photomicrographs from the outlined area from **(a)** show that almost all GFP-ir neurons **(b)** express *Vgat* mRNA hybridization signal **(c)** in the merged-channel image **(d)**. *Filled white arrowheads* indicate representative *Vgat*-ISH and GFP-ir colocalized neurons while *open arrowheads* point to a GFP-ir neuron that is vGAT-negative. Scale bars: 200 μm **(a)**; 25 μm **(b–d)**.

### 3.2: Distribution of GABAergic TH-ir neurons in the hypothalamus

The hypothalamus includes gray matter regions starting rostrally at the level of the anteroventral periventricular nucleus (AVP) and proceeding caudally until the emergence of the ventral tegmental area. In the mouse, the hypothalamus extends approximately 3 mm along the anteroposterior axis (+0.445 to –2.48 mm from Bregma), and this distance is represented across 33 levels (L; L50–L83) in the reference space of the *ARA* (Dong, 2008). We observed TH-ir neurons throughout the entire rostrocaudal extent of the hypothalamus, where they were dispersed in numerous gray matter regions. We evaluated the distributions of TH-ir neurons across the *ARA* reference space (Fig. 4a), including those TH-ir neurons that coexpress EGFP-f (Fig. 4b). Overall, there was a strong linear relationship between the total number of TH-ir neurons and the number of TH-ir neurons coexpressing EGFP-f (*r* = 0.969, *R^2^* = 0.939; Fig. 4c).

**Figure 4.**
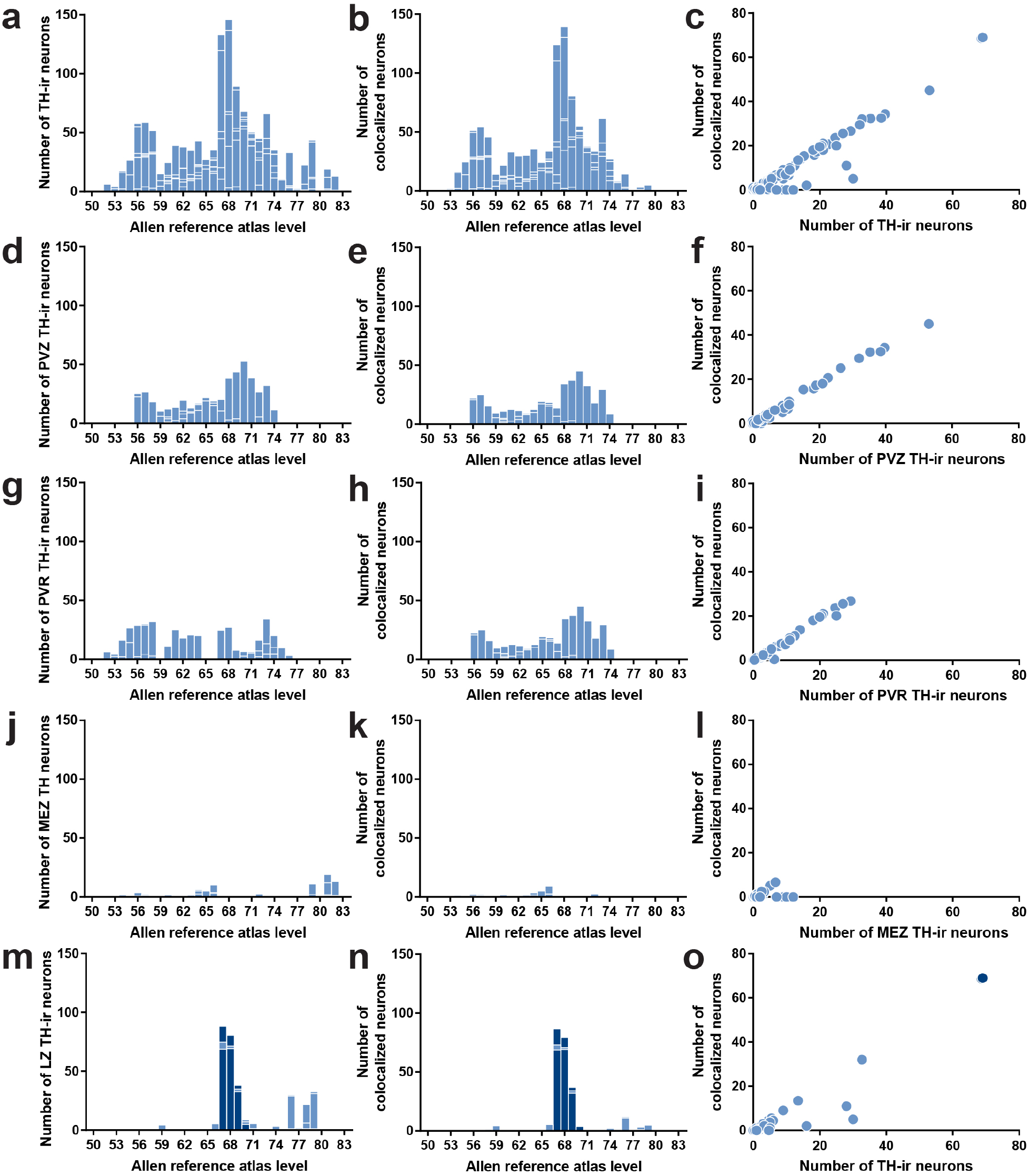
Distributions of TH-ir neurons with the colocalization of EGFP fluorescence in *Vgat-cre;L10-Egfp* brain tissue. Number of TH-ir neurons **(a, d, g, j, m)** that coexpress native EGFP fluorescence **(b, e, h, k, n)** at each atlas level throughout the whole hypothalamus **(a, b)**, periventricular hypothalamic zone (PVZ; **d, e**), periventricular hypothalamic region (PVR; **g, h**), hypothalamic medial zone (MEZ; **j, k**), and hypothalamic lateral zone (LZ; **m, n**). Correlation between the coexpression of native EGFP fluorescence in TH-ir neurons at all brain regions in each atlas level of the whole hypothalamus **(c)**, PVZ **(f)**, PVR **(i)**, MEZ **(l)**, and LZ **(o)**. *Dark blue bars* in **m** and **n** and the *dark blue filled circles* in **o** correspond to regions within the zona incerta.

In order to increase the spatial resolution for visualizing the distribution of TH-ir neurons in the hypothalamus, we mapped their locations onto *ARA* atlas templates that are available as electronic vector-onject files (Dong, 2008) (Fig. 5) and quantified the proportion of TH-ir neurons that coexpress EGFP-f in each parcellated gray matter region of the hypothalamus (Table 3). All regions with detectable labeling of TH-ir neurons are tabulated using the hierarchical organization of brain structures presented by the *ARA* (Allen Institute for Brain Science, 2011b). The colocalization of TH immunoreactivity with EGFP-f occurred at low (<50%), moderate (50–80%), or high frequencies (>80%) of colocalization.

**Table 3.**
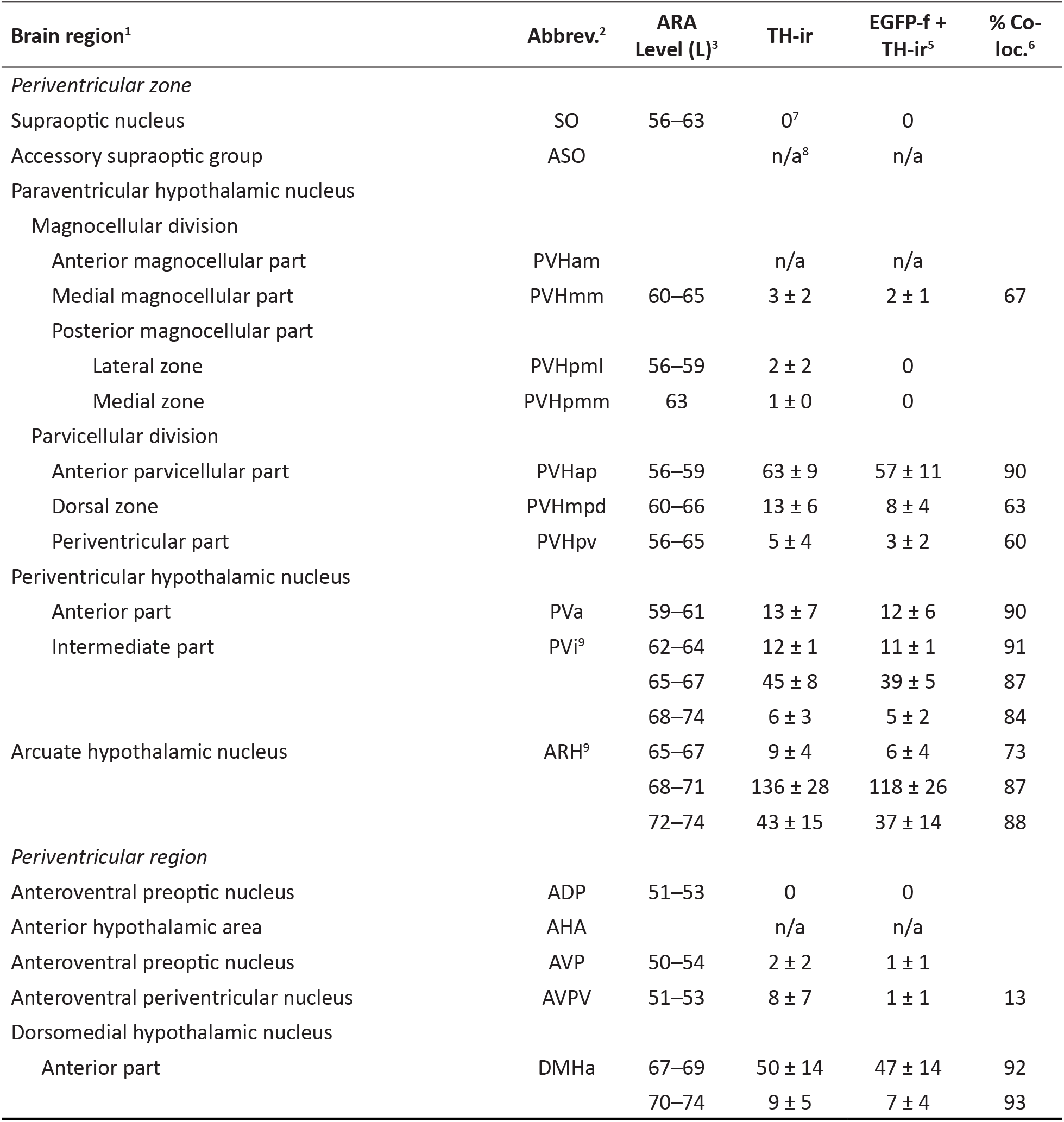
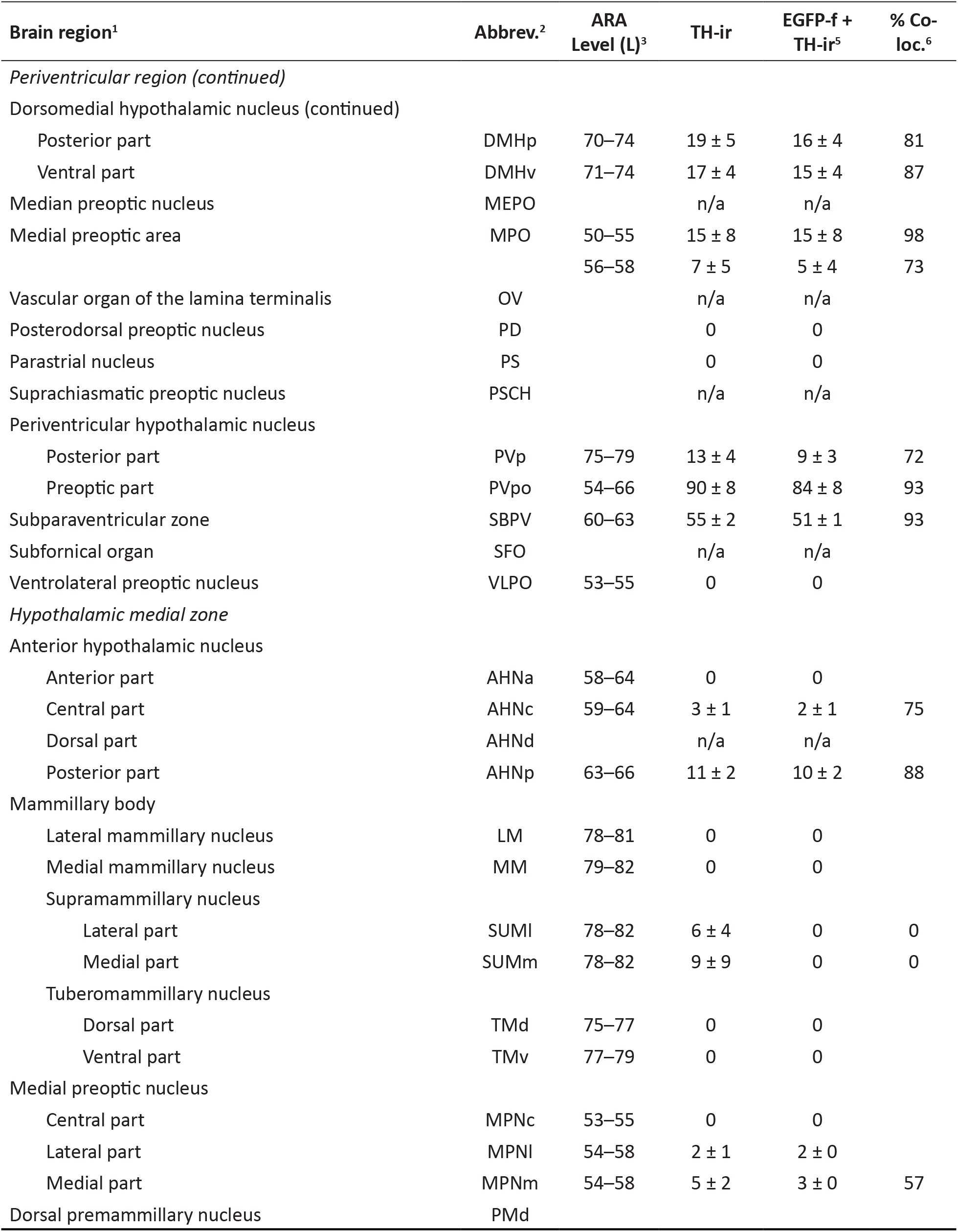
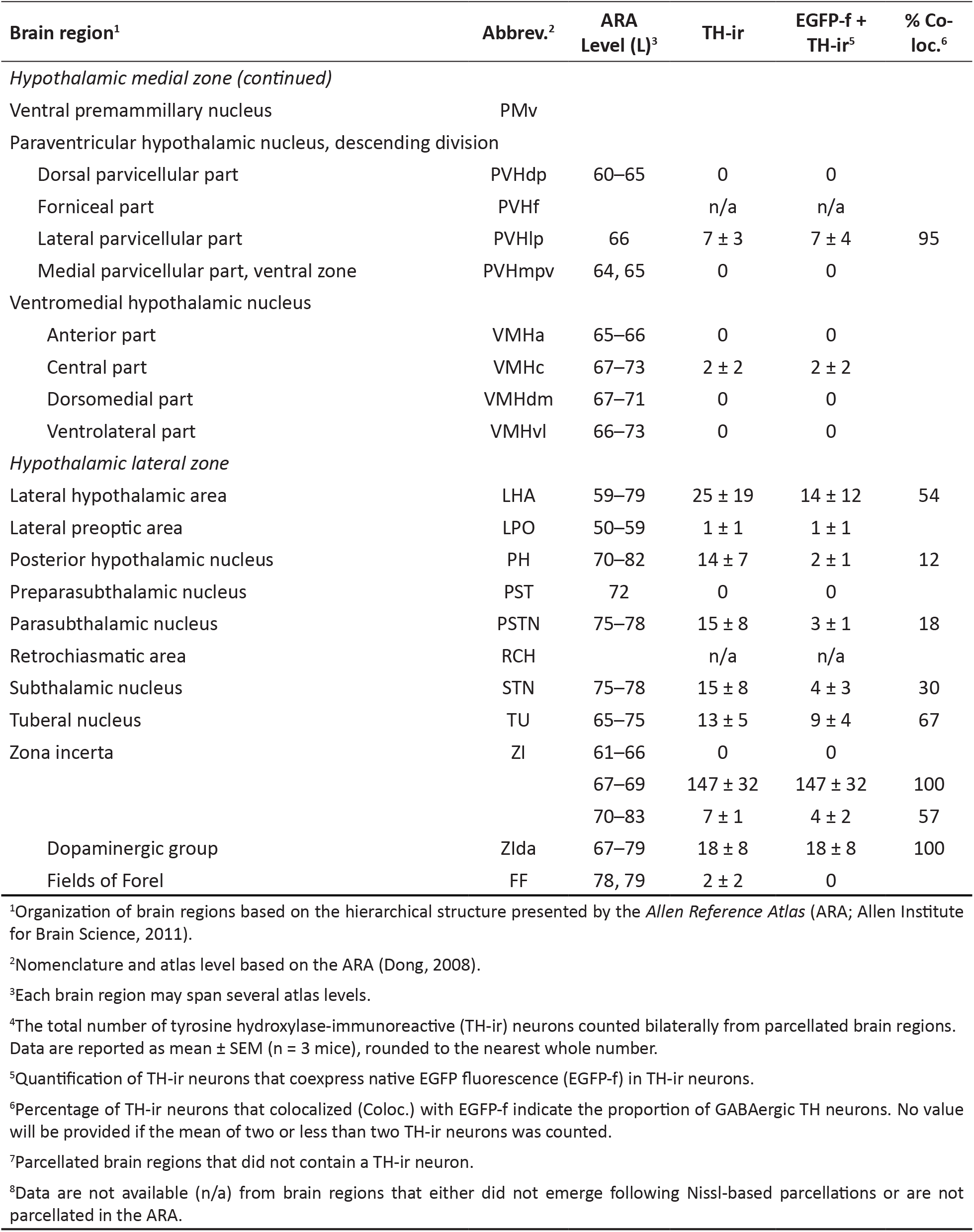
Quantification of EGFP-labeled TH-immunoreactive neurons from *Vgat-cre;L10-Egfp* mouse brain tissue

**Figure 5.**
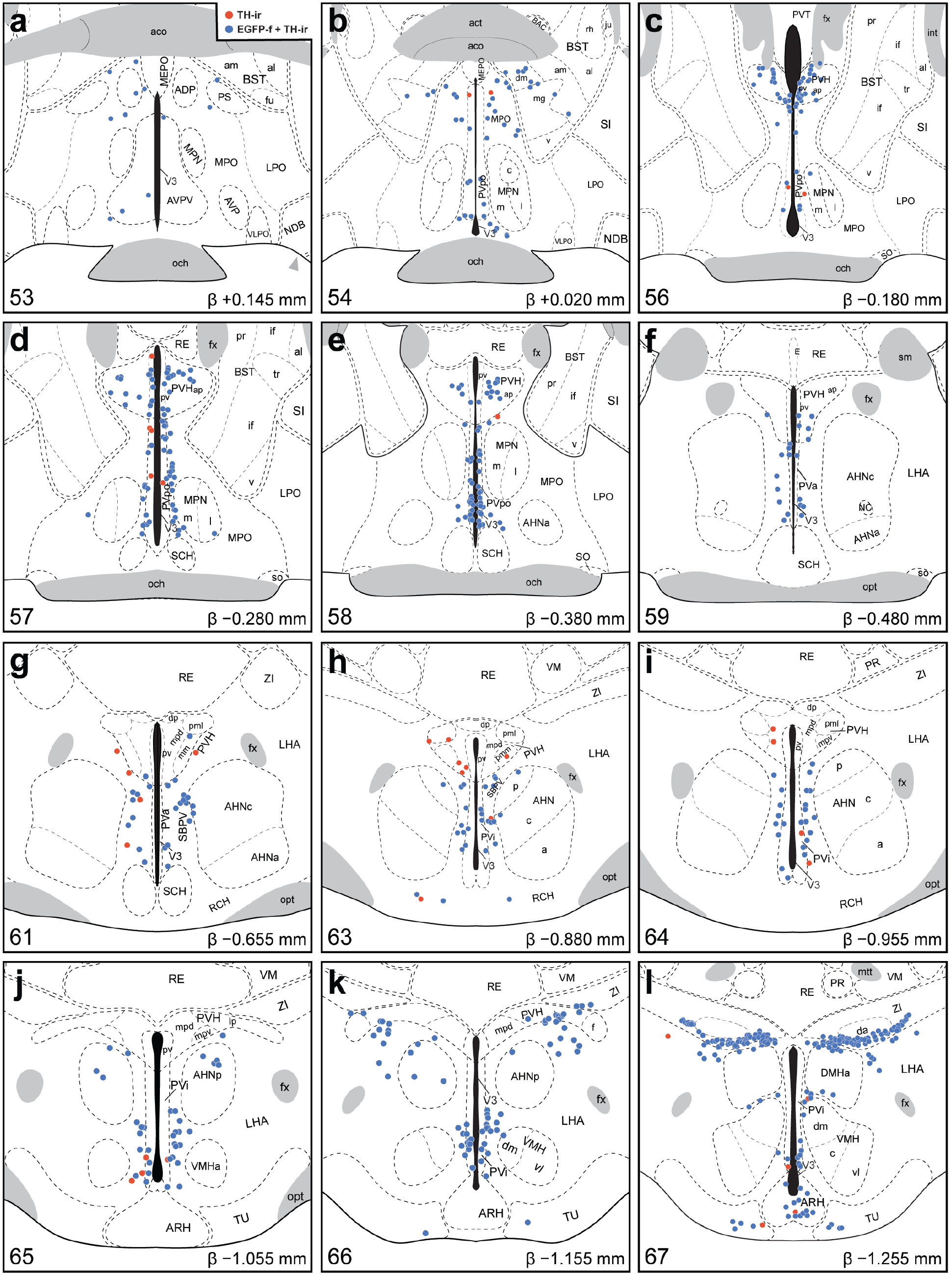
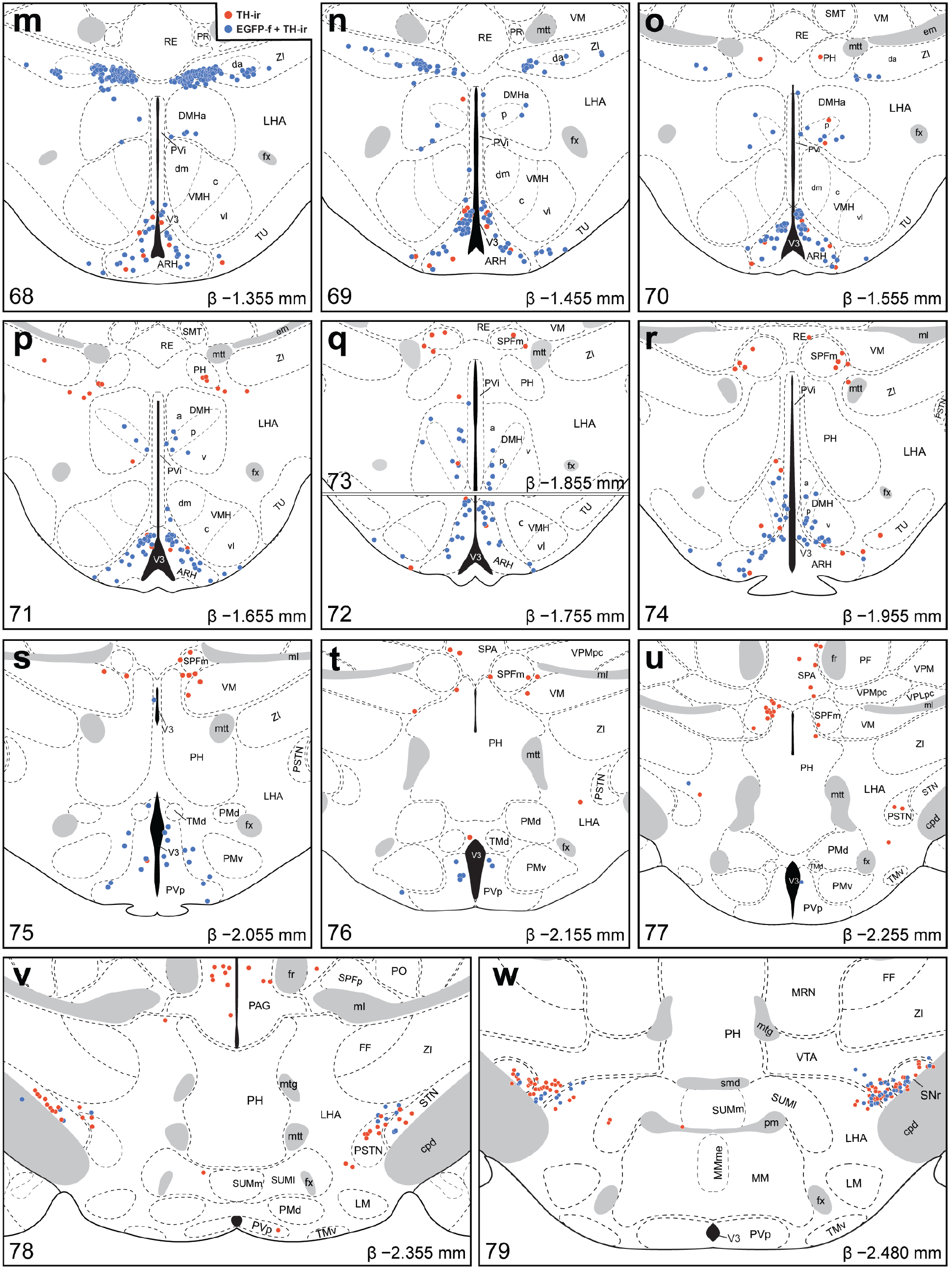
Mapped distributions of GABAergic TH-ir neurons in the hypothalamus. Representative transverse maps, arranged from rostral to caudal order **(a–w)**, show the distributions of TH-ir neurons that colocalize (*blue circles*) or do not colocalize (*red circles*) with native EGFP fluorescence from *Vgat-cre;L10-Egfp* mice. Consultation of Nissl-stained tissue guided the assignment of neurons to parcellated cytoarchitectonic boundaries in the TH-ir tissue, and plane-of-section analysis facilitated their mapping to gray matter regions of the *ARA* reference atlas templates. Each panel includes a portion of the atlas template for a given *ARA* level, the numerical designation of the atlas level (*bottom left*), the corresponding inferred stereotaxic coordinate from Bregma (β, in mm; *bottom right*), and the brain region labels adopted from formal nomenclature of the *ARA* (Dong, 2008).

#### 3.2.1: The periventricular zone

The TH-ir neurons in the periventricular zone were distributed within three main brain regions: the paraventricular hypothalamic nucleus (PVH), periventricular hypothalamic nucleus (PV), and arcuate hypothalamic nucleus (ARH) (Table 3). These neurons were distributed across L56–L74 and were moderately to highly (63–91%) colocalized with EGFP-f, which showed a very strong linear relationship with the overall numbers of TH-ir neurons in this zone (*r* = 0.996, *R^2^* = 0.992; Fig. 4d–f).

TH-ir neurons in the PVH were largely confined to its parvicellular division and were most numerous within the anterior parvicellular part (PVHap), and 90% of PVHap TH-ir neurons expressed EGFP-f (Table 3; L56–L59; Fig. 5c–f). In contrast, the dorsal zone (PVHmpd; Table 3) or periventricular part (PVHpv; L56–59; Fig. 5c–f; Table 3) of the parvicellular PVH contained much fewer TH-ir neurons, as 60% and 63% of PVHmpd and PVHpv neurons, respectively, also expressed EGFP-f (Table 3).

Most of the TH-ir neurons in the periventricular zone abutted the third ventricle. Within this zone, the entire periventricular hypothalamic nucleus that was mapped extended through 15 atlas levels, including the anterior (PVa) and intermediate (PVi) parts, and 84–90% of TH-ir neurons in this region expressed EGFP-f (L59–74; Fig. 5f–r; Table 3). Notably, the middle anteroposterior portion of the PVi (L65–L67) expressed the most EGFP- and TH-positive neurons (Fig. 5j–l). In the ARH, TH-ir neurons were distributed throughout the entire rostrocaudal extent of the structure, but were most abundant within the dorsomedial aspect of the ARH at L68–L71 (Fig. 5m–p). TH-ir neurons were distributed in varying densities through the ARH along its anteroposterior expanse, and 73–88% of these neurons colocalized EGFP-f (Table 3).

#### 3.2.2: The periventricular region

The periventricular region included brain regions across virtually the entire rostrocaudal extent of the hypothalamus, though TH-ir neurons were clustered either toward its rostral (L52–L63) or caudal portions (L67–L76) (Fig. 4g–h). Overall, most TH-ir neurons in the periventricular region also colocalized with EGFP-f (*r* = 0.993, *R*^*2*^ = 0.985; Fig. 4i), but we found regional differences at several atlas levels.

Within the rostral periventricular region, TH-ir neurons were most abundant in the preoptic part of the periventricular hypothalamic nucleus (PVpo; L54–58), and 93% of these neurons colocalized with EGFP-f (Table 3; Fig. 5b–e). Colocalized neurons that fell on or immediately outside the parcellated boundaries of PVpo in L57 were included in the cell counts for PVpo (Fig. 5d). There were also notable TH-ir neurons in the subparaventricular zone (SBPV; L60–63) that displayed very high colocalization with EGFP-f (Table 3; Fig. 5g, h). Other areas in the periventricular hypothalamic region contained a few TH-ir neurons in the anteroventral periventricular nucleus (AVPV) and medial preoptic area (MPO). Colocalization of EGFP-f in TH-ir neurons of the AVPV was low (13%; Table 3; Fig. 5a) but moderate to high (73–98%) in the MPO (Table 3; Fig. 5b–e).

Within the caudal periventricular region, TH-ir neurons were found in the posterior part of the periventricular hypothalamic nucleus (PVp; L75–79), where 72% of them colocalized with EGFP-f (Table 3; Fig. 5s–w). In the dorsomedial nucleus of the hypothalamus (DMH; Table 3), aside from a small cluster of cells lining the dorsomedial tip of its anterior part at L67 (Fig. 5l), TH-ir neurons were sparsely distributed throughout the remaining DMHa (L68–74), as well as the ventral (DMHv; L71–74), and posterior (DMHp; L70–74) parts of the DMH. TH-ir neurons throughout the entire DMH showed between 80–90% colocalization with EGFP-f (Table 3).

#### 3.2.3: The hypothalamic medial zone

The hypothalamic medial zone rarely contained TH-ir neurons (Fig. 4j, k), and overall, there was only a weak correlation between EGFP-f and TH-ir neurons (*r* = 0.402, *R^2^* = 0.162; Fig. 4l). There were few, if any, TH-ir neurons within the anterior (AHNa) or central (AHNc) parts of the anterior hypothalamic nucleus, but the posterior part (AHNp) contained distinct TH-ir neurons that colocalized with EGFP-f (Table 3; Fig. 5f–k). A few TH-ir neurons resided more caudally in the supramammillary nucleus (SUM) between L78–L82, but none of these neurons colocalized with EGFP-f (Table 3; Fig. 5v, w).

Notably, the PVH, which contained a moderate density of colocalized EGFP-f and TH-ir neurons within the periventricular zone, only displayed few such neurons within the hypothalamic medial zone (in its descending division), specifically in the lateral parvicellular part (PVHlp) (L66; Fig. 5k).

#### 3.2.4: The lateral hypothalamic zone

With the exception of the zona incerta (ZI), which accounted for most of the positive lateral hypothalamic zone TH-ir neurons between L66–69, the brain regions in this zone primarily contained a low density of TH-ir neurons (Fig. 4m), but there was a strong correlation between EGFP-f and TH-ir neurons (*r* = 0.952, *R^2^* = 0.905). In the anterior region of the lateral hypothalamic zone, the lateral hypothalamic and lateral preoptic areas contained few, if any, TH-ir neurons that colocalized EGFP-f. Posterior levels of the hypothalamus (L70–82), which include the posterior hypothalamic nucleus (PH), parasubthalamic nucleus (PSTN), subthalamic nucleus (STN), and tuberal nucleus (TU) contained few TH-ir neurons. With the exception of the TU TH-ir neurons, which had moderate (67%) colocalization (Table 3), the other regions displayed low (15–30%) colocalization with EGFP-f (Table 3).

In contrast, we found a high density of TH-ir neurons in the ZI, particularly within L67–L69 (Fig. 5l–n), and every TH-ir neuron in this space displayed EGFP-f (Fig. 5k–n). The majority of these TH-ir EGFP-f neurons lined the ventral edge of the ZI at L67 and clustered within the medial aspect of the ZI at L68. Interestingly, the majority of these neurons resided outside the cytoarchitectural boundaries of a cell group labeled in *ARA* reference atlas templates (Dong, 2008) as the dopaminergic group within the ZI (ZIda) in the ARA, although those that fell within its boundaries also displayed EGFP-f (Fig. 5l–n).

### 3.3: Striking distribution pattern of GABAergic TH-ir neurons within the zona incerta

While several hypothalamic regions displayed strong colocalization between TH immunoreactivity and EGFP-f, the distribution pattern of neurons with such colocalization was the most prominent and specific within the ZI. The ZI is a relatively expansive region that spans 2.33 mm along the rostrocaudal axis of the mouse brain, as represented within the *ARA* reference space at L61–L83. EGFP-f colocalized with all TH-ir ZI neurons found from L67 to L69 (Fig. 5l–n). The ZI was notable among brain regions because the degree of colocalization of TH-ir neurons with EGFP-f was the most complete among all brain regions analyzed at each atlas level (see right- and upper-most data points in Fig. 4, **panels c and o**). In contrast, there were very few TH-ir neurons anterior to L67 that were within the cytoarchitectonic boundaries of the ZI (Fig. 5g–k), only a few TH-ir ZI neurons that were located between L70–L83 (*see* Fig. 5o–w for L70–79), and only 57% of ZI neurons residing outside L67–L69 colocalized with EGFP-f (Table 3).

### 3.4: Neurochemical characterization of GABAergic TH-ir neurons

The hypothalamus is enriched by the expression of diverse neuropeptides, including those residing within or surrounding the ZI within the lateral hypothalamic zone, such as MCH and H/O. Previous reports have determined that MCH (Chee et al., 2015) and H/O neurons (Schöne et al., 2012) are glutamatergic. To confirm that the ZI TH-ir neurons do not colocalize these neuropeptides, we performed dual-label IHC reactions for TH and either MCH or H/O. Consistent with previous work and our findings that all TH-ir neurons in the ZI colocalize with EGFP-f, which indicates vGAT expression (Fig. 2), these reactions did not show the colocalization of either MCH (Fig. 6a–d) or H/O (Fig. 6e–h) immunoreactivity with TH-ir neurons.

**Figure 6.**
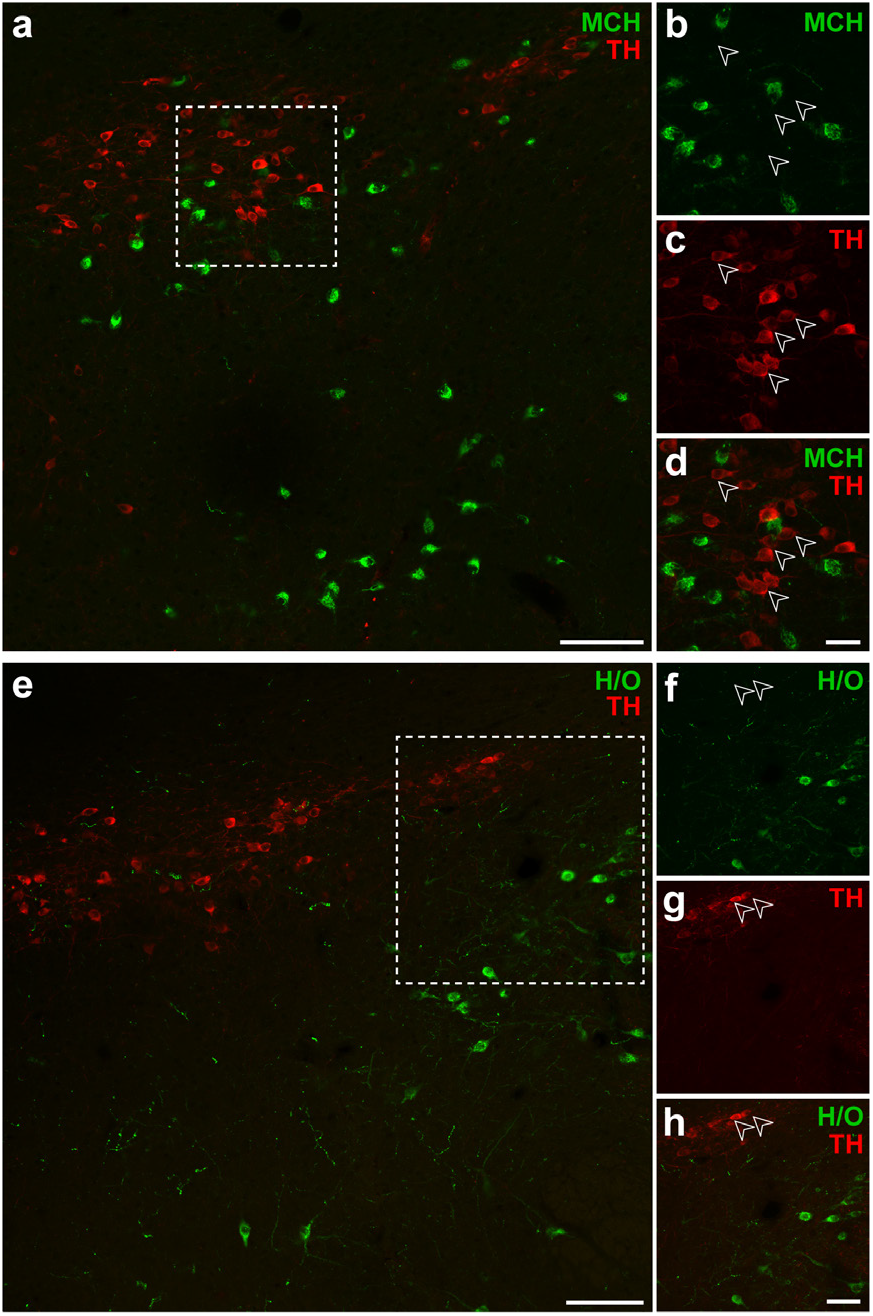
TH-ir neurons in the ZI do not express MCH or H/O. Confocal photomicrographs from the lateral hypothalamic zone of wild type mouse tissue illustrating the distributions of MCH-ir **(a–d)** and H/O-ir neurons **(e–h)** relative to TH-ir neurons. *White open arrowheads* show lack of MCH-(**b**) or H/O-positive labeling **(f)** in TH-ir neurons **(c, g)** in merged-channel, high-magnification photomicrographs **(d, h)** from the outlined areas in **a** and **e**, respectively. Scale bars: 100 µm **(a, e)**; 30 µm **(b–d)**; 50 µm **(f–h)**.

TH is the rate-limiting enzyme for all catecholamine neurotransmitter synthesis (Udenfriend & Wyngaarden, 1956) in mammals and is commonly used as a marker of catecholaminergic neurons. However, it does not further specify the terminal catecholamine (*i.e.*, dopamine, norepinephrine, or epinephrine) synthesized by the neuron. Within the catecholamine biosynthesis pathway, dopamine is hydroxylated by dopamine β-hydroxylase (DBH; EC 1.14.17.1) into norepinephrine which, in turn, can be methylated to become epinephrine. We did not observe DBH-ir neurons in any region within the hypothalamus, as dual-label IHC for DBH and TH immunoreactivities in *Vgat-cre;L10-Egfp* tissue did not reveal DBH-ir neurons within the ZI (Fig. 7). These results show that TH-ir ZI neurons lack the enzymatic machinery to synthesize norepinephrine, and hence also epinephrine. Therefore, we determined if TH-ir ZI neurons would display immunoreactivity to dopamine. To this end, we performed triple-label IHC for dopamine, TH, and GFP in the ZI of *Vgat-cre;L10-Egfp* mouse brain tissue and found that almost all GFP- and TH-ir neurons also displayed dopamine immunoreactivity (Fig. 8).

**Figure 7.**
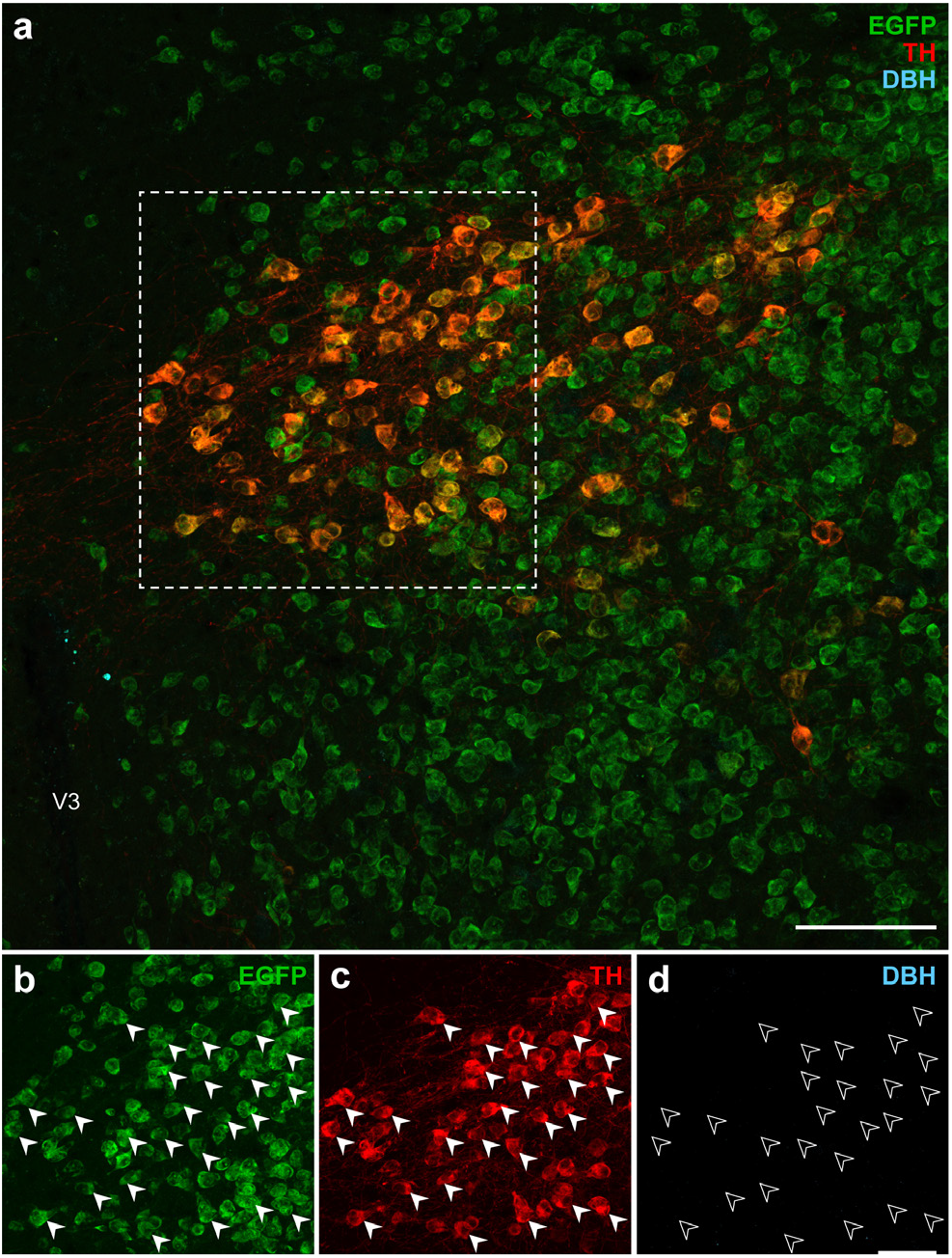
GABAergic TH-ir neurons in the ZI do not express dopamine β-hydroxylase. Confocal photomicrographs from the ZI of *Vgat-cre;L10-Egfp* brain tissue **(a)** show native EGFP fluorescence **(b)** in TH-ir neurons **(c)** but not immunoreactivity for dopamine β-hydroxylase (DBH; **d**). *Filled white arrowheads* mark a representative sample of GABAergic TH-ir neurons **(b, c)** from the outlined area **(a)** that do not express DBH, as indicated by *open white arrowheads* **(d)**. Scale bars: 100 µm **(a)**; 50 µm **(b–d)**. V3, third ventricle.

**Figure 8.**
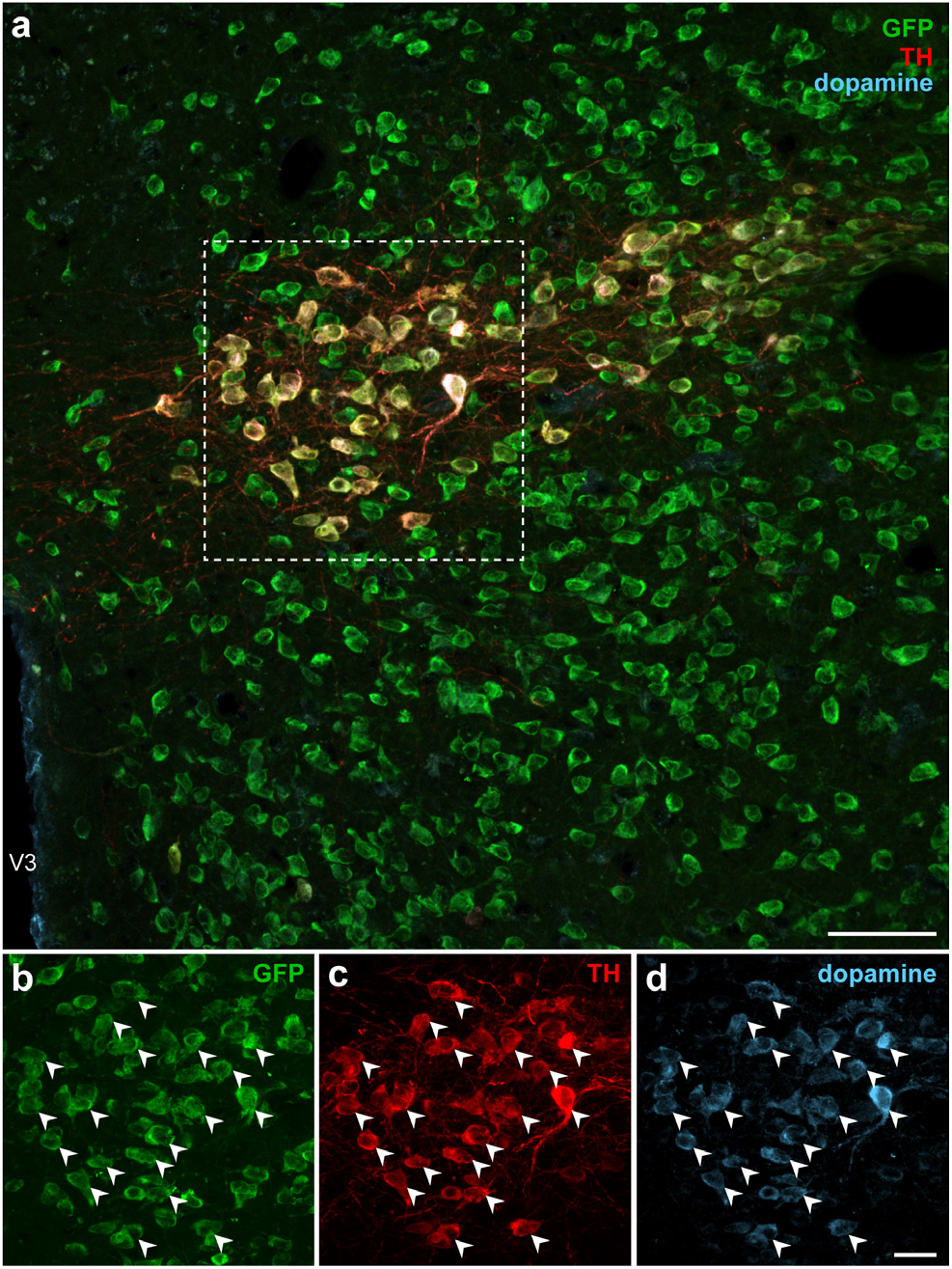
GABAergic TH-ir neurons in the ZI contain dopamine. Confocal photomicrographs from the ZI of *Vgat-cre;L10-Egfp* brain tissue **(a)** show the colocalization of GFP **(b)**, TH **(c)**, and dopamine **(d)** immunoreactivity at high magnification within the same neuron. *White filled arrowheads* indicate representative GFP-, TH-, and dopamine-ir neurons. Scale bars: 50 µm **(a)**; 20 µm **(b–d)**. V3, third ventricle.

## 4. Discussion

Hypothalamic catecholaminergic neurons can be marked by the expression of TH, the rate-limiting enzyme for catecholamine biosynthesis, and the majority of hypothalamic TH neurons also coexpress vGAT, and therefore may also signal via the neurotransmitter GABA. We mapped the locations of TH-ir neurons onto atlas templates from the *ARA* (Dong, 2008) to provide high-spatial resolution maps of their distribution throughout the hypothalamus. Several hypothalamic regions, including the arcuate and paraventricular hypothalamic nuclei, contained a high density of GABAergic TH neurons, but the most striking pattern of GABAergic TH neurons was observed within a highly circumscribed portion of the rostral ZI. We did not find that these ZI neurons colocalized with the few neuropeptide immunoreactivities (*i.e.*, MCH, H/O) that we examined; however, our neurochemical characterization to identify their catecholamine content revealed that these neurons also contain dopamine.

The use of a standardized neuroanatomical framework to visualize and map the distribution of TH-ir neurons onto atlas templates revealed their remarkably specific localization in a discrete portion of the ZI. The ZI spans more than 2.3 mm rostrocaudally, but almost all TH-ir neurons were found to reside specifically within a 0.2 mm-long segment of the ZI between L67–L69. Furthermore, these TH neurons appear to be homogeneous in their neurochemical makeup based on their colocalization of vGAT and dopamine, but not MCH or H/O neuropeptides, or DBH, which would permit additional downstream catecholamine biosynthesis (*e.g.*, norepinephrine and epinephrine). Together, the evidence in the present study indicates that these neurons likely produce dopamine as their terminal catecholamine in this biosynthetic pathway.

Our Nissl stains in the mouse brain highlight the preence of a compact region within the ZI, which was also labeled ZIda according to *ARA* nomenclature (Dong, 2008), but it is notable that the observed GABAergic TH neurons in our study were medial and ventral to this compact cell group. It has been previously reported that the medial and ventral/lateral distributions of TH neurons in the ZI may be differentiated by cell diameter (Ruggiero et al., 1984), but we only considered cell location and not cell size in our observations. Commonly used mouse brain atlases (Dong, 2008; Franklin & Paxinos, 2012) make reference to the A13 or dopaminergic cell group of the ZI, respectively, based on cytoarchitectural features. However, given that GABAergic TH neurons in the mouse ZI fall outside of the compact region assigned to these labels, the compact cell group in the mouse ZI may only approximate the location of dopaminergic neurons. It is noteworthy that the most recent iteration of the *ARA* nomenclature and online atlas levels (Allen Institute for Brain Science, 2011a) have removed the cytoarchitectonic boundaries originally assigned in the ZI to the compact A13 region. However, we have used *ARA* reference atlas templates (Dong, 2008), which still contain this region as a discrete structure (ZIda), to map ourTH-ir neuronal distributions – as these are at the time of this writing the only electroncally available vector-object templates for *ARA* reference space. As subsequent versions of the *ARA* become available, we hope that accompanying vector-object templates will be available for the community to use to map their datasets.

Transcriptomic analysis of neurons in the lateral hypothalamic area has also demonstrated the GABAergic and dopaminergic profile of TH neurons, which express genes encoding enzymes to synthesize GABA (*Gad1* and *Gad2*) and dopamine (*Ddc* and *Th*) as well as their respective vesicular transporters *Slc32al* (vGAT) for GABA and *Slc18a2* (vMAT2) for dopamine (Mickelsen et al., 2019). ZI TH neurons can synthesize dopamine and transport it into synaptic vesicles using the vesicular monoamine transporter, vMAT2 (Sharma et al., 2018). Similarly, ZI TH neurons also expresses glutamate decarboxylase (GAD) to synthesize GABA (Shin et al., 2007) and we showed here that all ZI TH neurons express vGAT to transport GABA into synaptic vesicles (Fig. 7, 8); thus they have the machinery to mediate GABAergic neurotransmission. By contrast, we did not find that ZI TH neurons colocalized with at least two major neuropeptides found in the lateral hypothalamic zone and, in light of the recent transcriptomic analysis of TH neurons in this area (Mickelsen et al., 2019), we also do not expect that ZI TH neurons would colocalize with another neuropeptide. MCH and H/O neurons do not overlap in the same cytoarchitectural space and both are glutamatergic (Chee et al., 2015; Mickelsen et al., 2019). Moreover, TH neurons do not show enriched gene expression for neuropeptides such as cocaine- and amphetamine-regulated transcript protein, galanin, neuropeptide W, neuropeptide Y, neurotensin, somatostatin, and tachykinin that are expressed in GABAergic neurons in the lateral hypothalamus (Mickelsen et al., 2019).

GAD expression was once commonly used to identify GABAergic neurons, but recent studies have shown that GABA synthesis in neurons does not necessarily signify that such neurons also release it (Williams et al., 2014; Chee et al., 2015). We therefore chose to identify GABAergic neurons based on vGAT expression, which we visualized by cre-dependent EGFP expression in *Vgat-cre* neurons. Gross, qualitative comparison of *Vgat* hybridization signals had shown its expression in expected brain regions (Vong et al., 2011). Therefore, we performed quantitative analyses to address any possible ectopic EGFP expression at the neuronal level. At least in the hypothalamus, we found that more than 99% of EGFP-expressing neurons expressed *Vgat* mRNA, thus confirming robust and accurate labeling of vGAT neurons in the hypothalamus of *Vgat-cre* mice. GABA is often reported as a co-transmitter in dopaminergic cells. For example, dopaminergic neurons in the ventral tegmental area may transport GABA and dopamine via the same vesicular transporter, vMAT2, suggesting that these two chemical messengers may be co-packaged (Tritsch et al., 2012). However, in other regions such as the olfactory bulb, GABA and dopamine are not necessarily co-packaged, and they may utilize independent modes of vesicular release that occur over different time courses (Borisovska et al., 2013). We are pursuing histological studies to identify the projection targets of ZI TH neurons (Mejia et al., 2019), as well as functional studies to determine if these purportedly GABAergic and dopaminergic ZI neurons do in fact release GABA and/or dopamine at downstream neurons.

GABAergic neurons within the ZI have recently been shown to modulate the motivational and hedonic aspects of feeding through direct projections to the paraventricular nucleus of the thalamus (PVT) (Zhang & van den Pol, 2017), but it is not known if dopamine also acts at the PVT to stimulate feeding. The PVT receives projections from ZI TH neurons (Li et al., 2014) and expresses dopamine D2 receptors, which are linked to its role in drug addiction (Clark et al., 2017). There is at least a foundational basis for dopamine- and GABA-mediated transmission in the PVT, but it remains to be determined if these chemical messengers would mediate the same behavioral roles at the PVT or elsewhere. For instance, another ZI GABA pathway targets the periaqueductal gray (PAG), and is implicated in the control of flight and freezing responses (Chou et al., 2018). A neuroanatomical tracing study showed ZI TH-ir projections to the PAG in rats (Messanvi et al., 2013) allowing for the possibility of a role for dopamine in mediating defensive behaviors.

Our work broadly supports the view that GABA co-transmission is a ubiquitous feature of dopaminergic neurons (Tritsch et al., 2016). Overall, we found vGAT-EGFP signals colocalized in 82% of hypothalamic TH-ir neurons, though some regions showed higher or lower levels of colocalization (Table 3). Outside the ZI, the largest number vGAT-positive TH-ir neurons were distributed through the PVH, PV, and ARH, regions which harbor neurons that project to the pituitary (Ju et al., 1986) and mediate neurosecretory functions (Markakis & Swanson, 1997). For example, TH-expressing neurons send projections to the median eminence to enable dopamine-mediated inhibition of prolactin release from the anterior pituitary gland (Ben-Jonathan et al., 1977; Ben-Jonathan & Hnasko, 2001). Moreover, cells in the anterior pituitary express all GABA receptor subtypes (Anderson & Mitchell, 1986) and GABA administration can also inhibit prolactin release (Grandison & Guidotti, 1979). Thus, in addition to mediating local neurocircuitry to regulate behavior, colocalized GABA and dopamine in neurons could work synergistically to mediate neuroendocrine functions.

While hypothalamic TH neurons have been reported since the 1970s (Hökfelt et al., 1976), our maps provide high spatial-resolution analysis of TH-ir neurons in the mouse hypothalamus. Further, this dataset is enriched by defining and mapping the distribution of TH neuron subpopulations that may also be GABAergic. Importantly, these maps constitute a standardized dataset that can be used to derive precise stereotaxic coordinates to target these discrete neuronal subpopulations in functional studies.

## Acknowledgments

This work is supported by NSERC Discovery Grant RGPIN-2017-06272 (MJC), NIH grants GM109817 and GM127251 (AMK), UTEP Office of Research and Sponsored Projects (AMK), and NIH center grant (2U54MD007592) funds to AMK awarded to the Border Biomedical Research Center at UTEP.

## Notes

#### Summary of Updates

Corrections to text.

